# Ultrastructure of light-activated axons following optogenetic stimulation to produce late-phase long-term potentiation

**DOI:** 10.1101/799890

**Authors:** Masaaki Kuwajima, Olga I. Ostrovskaya, Guan Cao, Seth A. Weisberg, Kristen M. Harris, Boris V. Zemelman

## Abstract

Analysis of neuronal compartments has revealed many state-dependent changes in geometry but establishing synapse-specific mechanisms at the nanoscale has proven elusive. We co-expressed channelrhodopsin2-GFP and mAPEX2 in a subset of hippocampal CA3 neurons and used trains of light to induce late-phase long-term potentiation (L-LTP) in area CA1. L-LTP was shown to be specific to the labeled axons by severing CA3 inputs, which prevented back-propagating recruitment of unlabeled axons. Membrane-associated mAPEX2 tolerated microwave-enhanced chemical fixation and drove tyramide signal amplification to deposit Alexa Fluor dyes in the light-activated axons. Subsequent post-embedding immunogold labeling resulted in outstanding ultrastructure and clear distinctions between labeled (activated), and unlabeled axons without obscuring subcellular organelles. The gold-labeled axons in potentiated slices were reconstructed through serial section electron microscopy; presynaptic vesicles and other constituents could be quantified unambiguously. The genetic specification, reliable physiology, and compatibility with established methods for ultrastructural preservation make this an ideal approach to link synapse ultrastructure and function in intact circuits.

## Introduction

The cellular correlates of learning and memory have been the subjects of intense study and speculation. We and others have used patterns of activity that produce late-phase long-term potentiation (L-LTP), a form of synaptic plasticity resulting in an increased synaptic efficacy. L-LTP is protein synthesis dependent and lasts more than three hours. Although some structural changes occur early following the induction of LTP, the lasting changes are most likely to reflect mechanisms of memory. *Post hoc* three-dimensional reconstruction from serial section electron microscopy (3DEM) of synapses and resident structures has revealed alterations that occur and are sustained long after the induction of LTP [1–4]. Mechanistic interpretation has been limited, however, because it has only been possible to compare subpopulations of synapses near LTP-producing versus control sites, rather than identify potentiated synapses. Genetic targeting to tag and stimulate individual cells can be achieved by co-expressing channelrhodopsin2 (ChR2) and a modified ascorbate peroxidase [5,6]. Here, we adapted this approach to potentiate a subset of CA3→CA1 hippocampal axons and to identify synapses recently involved in L-LTP.

Optical modulation of plasticity is routinely used across brain areas, and *in vivo* experiments have revealed a correlation between behavioral memory and optically-induced synaptic plasticity (LTP and long-term depression [LTD]) [7,8]. In slice preparations, past efforts using various protocols to induce LTP using ChR2 have been limited to whole-cell recordings and short post-induction times [9–12]. In the hippocampal CA3→CA1 pathway, a previous study used optical stimulation to induce late-phase LTD *in vivo* [13]; however, in this pathway, optically induced L-LTP to our knowledge has not been demonstrated.

We expressed ChR2 and labeled a subset of CA3→CA1 Schaffer collateral and commissural fiber axons using a single virus and then produced L-LTP using high-frequency light pulses. This approach generated a mosaic of labeled and unlabeled axons and allowed a within-preparation comparison of identified synapses having distinct histories of activation. As proof of concept, we reconstructed a labeled axon through 3DEM from a slice that displayed optically induced L-LTP. The tissue quality was superb, demonstrating that we could identify genetically tagged activated axons. The approach proved compatible with conventional tissue fixation and the processing methods needed to preserve subcellular organelles. Hence, it provides a reliable strategy to study synapse-specific mechanisms of synaptic plasticity.

## Materials and methods

### Animals

This study was carried out in accordance with the recommendations in the Guide for the Care and Use of Laboratory Animals of the National Institutes of Health. All animal procedures were approved by the University of Texas at Austin Animal Care and Use Committee (protocol number AUP-2012-00056 and its successors). All mice were housed under reversed light/dark cycles in an AAALAC-accredited facility managed by the University of Texas Animal Resource Center. We used 8-12 week old male 129S6/SvEvTac mice (Taconic Biosciences, RRID:IMSR_TAC:129sve) for all LTP experiments. Male C57B/6J mice (The Jackson Laboratory, RRID:IMSR_JAX:000664) were also used for earlier experiments, which are indicated as such in figure captions where applicable. All efforts were made to minimize suffering.

### AAV assembly and production

A channelrhodopsin2^ET/TC^ [14] fusion protein was assembled with superfolder green fluorescent protein (GFP) [15] fitted with C-terminal Kir2.1 ER export signal [16]. To generate mAPEX2, we modified the wild type ascorbate peroxidase from *P. sativum* (APX) [17] to include an N-terminal palmitoylation tag from growth-associated protein 43 (GAP-43) [18], amino acid substitutions K14D, W41F, E112K [5], and A134P [6], and a C-terminal hemagglutinin epitope tag (HA tag: YPYDVPDYA). ChR2 and codon-optimized mAPEX2 were separated by the self-cleaving porcine teschovirus P2A peptide [19] to produce both proteins from a single transcript. In earlier experiments, we used a version of the rAAV that encoded myc-tagged mAPEX1 with the first three mutations instead of mAPEX2 (indicated in figure captions where applicable).

In addition to the two proteins, the recombinant adeno-associated virus (rAAV) construct comprised an enhanced human synapsin promoter [20], the woodchuck post-transcriptional regulatory element and SV40 polyadenylation sequence. Viruses were assembled using a modified helper-free system (Stratagene) as serotype 2/1 (*rep/cap*) and purified on sequential cesium gradients according to published methods [21]. Titers were measured using a payload-independent qPCR technique [22]. Typical titers were >10^10^ viral genomes/µl.

### Rat hippocampal neurons

Hippocampal neurons obtained from rats (embryonic day 19) were grown in dissociated cultures [23] on coverslips and were infected on day 8 after plating with the rAAV construct encoding mAPEX1. At 6-10 days post-infection, the neurons used for immunostaining were fixed with 4% formaldehyde in phosphate-buffered saline (PBS) for 15 min. For labeling with 3,3′-diaminobenzidine (DAB; Sigma-Aldrich), the neurons were fixed for 30 min with 2% formaldehyde and 6% glutaraldehyde. All glutaraldehyde-containing fixative was prepared in 0.1 M sodium cacodylate buffer (pH = 7.4) with 2 mM CaCl_2_ and 4 mM MgSO_4_.

### Stereotaxic injections

Mice under isoflurane anesthesia (1-4% mixed in O_2_) were placed in a stereotaxic apparatus and prepared for injections with craniotomies over the hippocampal area CA3. Unilateral injections were performed using a pulled glass pipette (10-15 μm diameter tip) mounted on a Nanoject II small-volume injector (Drummond Scientific). Approximately 30 nl of virus was deposited at each injection site at 1-2 minute intervals (from bregma in mm: AP +1.9, ML −2.3, DV 1.8, 1.6, 1.4; AP +2.1, ML −2.5, DV 1.8, 1.6, 1.4). The pipette was left in place for 3–5 min before being removed from the brain. Carprofen (5 mg/kg, sc; TW Medical Cat# PF-8507) was injected 20 min before the end of surgery, and mice were monitored daily thereafter to ensure full recovery. Two rAAV-injected mice (129S6/SvEvTac) were perfusion-fixed under heavy isoflurane anesthesia with 4% paraformaldehyde in 0.02 M phosphate buffer (PB) to verify injection sites, and 15 mice (four 129S6/SvEvTac and 11 C57B/6J) were perfusion-fixed with glutaraldehyde (up to 2.5%) and formaldehyde (up to 2%), followed by 20 mM glycine in cacodylate buffer to quench free aldehydes, to verify enzymatic activity of mAPEX2 with Ni-DBA staining as described below.

### Histology and light microscopy

For immunostaining, the fixed neurons were permeabilized with 0.2% Triton X-100 in PBS for 5 min, rinsed in PBS, and blocked in 5% bovine serum albumin (BSA; Sigma-Aldrich) and 5% normal goat serum (NGS; VectorLabs) for 15 min. The cells were then incubated overnight at 4°C with rabbit anti-myc (1:250; Sigma-Aldrich Cat# C3956, RRID:AB_439680) in PBS with 2% BSA, 3% NGS, 0.1% Triton X-100, followed by PBS washes and incubation for 1 hr with goat anti-rabbit IgG conjugated with Cy5 (1:100; Jackson ImmunoResearch Labs Cat# 111-175-144, RRID:AB_2338013) in PBS with 2% BSA, 3% NGS, 0.1% Triton X-100. After PBS washes, the coverslips containing neurons were mounted on glass microscope slides with Aqua/Poly antifade mountant (PolyScience) for epifluorescence microscopy.

For DAB-labeling, the neurons were rinsed with cacodylate buffer and treated with 20 mM glycine. Then the neurons were rinsed several times before being incubated with DAB (0.5 mg/ml) and H_2_O_2_ (0.03%) in cacodylate buffer for 30 min. After buffer rinses, the coverslips were dehydrated in ethanol, cleared in xylenes, and mounted on glass slides with DPX (Electron Microscopy Sciences) for brightfield microscopy.

To verify injection sites, the perfusion-fixed brain was vibratome-sectioned (100 µm thickness) for epifluorescent microscopy to visualize GFP. To assess enzymatic activity of mAPEX2, the vibratome-sections (50 µm thickness) of the fixed brain containing the dorsal hippocampus were incubated with Ni-DAB solution (2.5 mM ammonium Ni [II] sulfate and 0.8 mM DAB in 0.1 M PB) for 20 min, before H_2_O_2_ (final concentration 0.0003%) was added and incubated for 10 min. After PB rinses, some of the Ni-DAB labeled sections were processed for EM as described below. Otherwise, they were mounted on glass microscope slides, dehydrated in ethanol, cleared with xylenes, and coverslips were applied with DPX. Epifluorescence and brightfield images were acquired on a Zeiss Axio Imager.Z2 microscope with AxioCamMR3 camera, or a Zeiss Axio Imager.M2 with AxioCamHRc3 camera.

Instead of the TSA labeling (described below), some of the vibratome sections collected from fixed hippocampal slices were permeabilized and blocked with PBS containing 0.3% Triton X-100, 1% BSA, and 10% NGS. The vibraslices were then incubated for overnight at RT with rabbit anti-HA antibody (1:1000; Cell Signaling Technology Cat# 3724, RRID:AB_1549585), followed by 1 hr with the Cy5-conjugated goat anti-rabbit IgG. After PBS rinses, the vibraslices were mounted on glass microscope slides and coverslips were applied with Aqua/Poly mountant for imaging with a Leica TCS SP5 confocal microscope. Single channel stacks (8 bit, 2048 × 2048 pixels at 19.1 or 28.6 nm/pixel) were acquired for GFP (458 nm laser) and Cy5 (633 nm laser) with 63× objective (oil, NA 1.32) at 4× zoom.

### Slice preparation, electrophysiology, optical stimulation

Six weeks after rAAV injections, the mice were anesthetized deeply with isoflurane and then decapitated. The brain was removed from the cranium, and the left hippocampus was dissected out and rinsed with room temperature (RT) artificial cerebrospinal fluid (aCSF; pH = 7.4) containing (in mM) 117 NaCl, 5.3 KCl, 26 NaHCO_3_, 1 NaH_2_PO_4_, 2.5 CaCl_2_, 1.3 MgSO_4_, and 10 D-glucose, and bubbled with 95% O_2_-5% CO_2_. Slices (400 μm thickness; 4 per animal) from the dorsal hippocampus were cut at 70° transverse to the long axis on a Stoelting tissue chopper and transferred in oxygenated aCSF to the supporting nets of interface chambers in the Synchroslice system (Lohmann Research Equipment). The entire dissection and slice preparation took ~5 min. This rapid dissection, together with the interface chamber design, provides for high-quality ultrastructure during long acute slice experiments [24–27]. Hippocampal slices were placed on a net at the liquid-gas interface between 30-31°C aCSF and humidified 95% O_2_-5% CO_2_ atmosphere bubbled through 35-36°C deionized water. For experiments in which the area CA3 was cut, the dissection was made using the 25-gauge needle on a 1-ml syringe after the slices were transferred into the chambers.

After 3 hr of incubation, the optical fiber (300 μm core diameter; 0.39 NA; ThorLabs FT300UMT) and recording electrode (Thomas Recording) were positioned 400-600 μm apart in middle *stratum radiatum* of the area CA1 with the fiber placed toward the area CA3. An optimal position of the recording electrode was chosen by probing several points across CA1 and SR to achieve larger responses. One-ms pulses of laser (λ = 473 nm; maximal output ~14.5 mW as measured by a Thorlabs S130C light meter) were delivered from source (ThorLabs S1FC473MM) controlled by a pulse generator (A.M.P. Instruments Master-8). The light meter placed directly under the hippocampal slices showed approximately 30% loss of laser power through the slice thickness, while the recording site was approximately 100 µm deep from the top surface. GluN receptors were blocked by adding 4 μl of 25 mM d,l-2-amino-5-phosphonovaleric acid (APV; Abcam Cat# ab144498) to the 1 ml of aCSF in the interface recording chamber, which achieved an effective concentration of 50 μM d-APV. Tetrodotoxin (TTX; final concentration 1 μM; Abcam Cat# ab120055) and 6,7-dinitroquinoxaline-2,3-dione (DNQX; final concentration 20 μM; Abcam Cat# ab144496) were added to block voltage-gated sodium channels and GluA receptors, respectively.

For electrical stimulation experiments, we replaced the optical fiber with a concentric bipolar electrode (FHC, Inc.), which was used to deliver 200 μs biphasic current pulses (100-300 μA), lasting 100 μs each for positive and negative components of the stimulus. The initial eEPSP slope was ∼50% of the maximal eEPSP slope based on the input-output (IO) curve for each slice. IO curves were recorded by using a sequence of pulses applied each 30 s with increasing stimulus intensity in 25 μA increments. Test pulses were given at 1 pulse per 2.5 min unless stated otherwise, and eEPSP was recorded. Paired-pulse ratio (PPR) was measured by applying two optical stimuli spaced 50-200 ms apart with 50 ms increment, every 30 s.

### Microwave-enhanced chemical fixation of hippocampal slices and TSA labeling

At end of recordings, the slices were immersed in fixative containing 1% glutaraldehyde and 4% formaldehyde, and were microwaved immediately for 8-10 s at 700 W (modified from ref. 27). The fixed slices were stored overnight at RT in the same fixative or in cacodylate buffer. After being immersed in 20 mM glycine for 20 min and buffer rinses, the area CA1 was dissected out under a stereoscope with a microknife and embedded into 9% agarose. Vibratome sections (50 µm thickness) were then collected from the area of optical stimulation to the location of the recording electrode. ChR2-GFP expression was confirmed with epifluorescence microscopy. The sections were transferred to 0.1 M PB and then incubated with tyramide-conjugated Alexa Fluor 647 (Thermo Fisher Scientific Cat# T20951) for 15 min in dark before H_2_O_2_ was added (final concentration 0.0015%) and incubated for additional 10 min in dark. After washes with PB, the vibraslices were washed in cacodylate buffer before being processed for 3DEM.

### Tissue processing for 3DEM

TSA-labeled vibratome section was embedded into 9% agarose to protect it during the subsequent processing, as described previously [28,29]. The tissue was immersed for 5 min in reduced osmium solution containing 1% osmium tetroxide (OsO_4_; Electron Microscopy Sciences) and 1.5% potassium ferrocyanide (Sigma-Aldrich) in cacodylate buffer. After several buffer rinses, the tissue was immersed in 1% OsO_4_ and two cycles of microwave irradiation (175 W; 1 min on → 1 min off → 1 min on) were applied with cooling to ~15°C in between. The tissue was rinsed in buffer several times, twice in purified water, and then immersed in 50% ethanol before being dehydrated in ascending concentrations of ethanol (50%, 70%, 90%, 100%) containing 1% uranyl acetate (UA; Electron Microscopy Sciences) with application of microwave irradiation (250 W, 40 s per ethanolic UA step). Ethanol was replaced by propylene oxide, and the tissue was infiltrated and embedded into LX-112 resin (Ladd Research). Embedded tissue was trimmed under a stereomicroscope to expose the region of interest containing the middle *stratum radiatum* of the area CA1. Serial thin sections (~60 nm thickness) were cut with a diamond knife (Diatome Ultra35) on a ultramicrotome (Leica Ultracut UC6 or UC7) and collected on Synaptek TEM grids (Be-Cu or gilded; Electron Microscopy Sciences or Ted Pella) coated with film of polyetherimide (PEI; Goodfellow).

### Post-embedding immunogold labeling and gold enhancement

Serial thin sections on gilded grids (4-6 sections per grid) were rinsed with Tris-buffered saline (TBS; pH = 7.6) containing 0.01% Triton X-100 (TBS-T) and then blocked with 2% human serum albumin (HSA; Sigma-Aldrich) and 10% NGS in TBS-T for 30 min. The sections were then incubated overnight at 4°C with TBS-T containing 1% HSA, 1% NGS, and mouse anti-Cy5/Alexa Fluor 647 (cocktail of antibodies at 1:100 each from Sigma-Aldrich [Cat# C1117, RRID:AB_477654] and Miltenyi Biotec [custom-ordered antibody used in their Anti-Cy5/Anti-Alexa Fluor 647 MicroBeads, Cat# 130-091-395]). After extensive washes with TBS-T and TBS (pH = 8.2; TBS-8.2), the sections were incubated for 90 min at RT with TBS-8.2 containing 1% NGS, 0.5% polyethylene glycol, and goat anti-mouse antibody conjugated with colloidal gold (5 or 15 nm diameter; BBI Solutions Cat# EM.GMHL5 or EM.GMHL15; 1:100). The sections were subsequently washed with TBS-8.2 containing additional 500 mM NaCl to reduce nonspecific antibody binding and then with TBS-8.2. The sections labeled with 5 nm gold were further rinsed with purified water, incubated with gold enhancement solution (GoldEnhance EM Plus; Nanoprobes) for 5 min at RT under ambient room light, and extensively washed in purified water. Shortening the incubation time for gold enhancement should reduce formation of background particles. All sections were stained with saturated aqueous solution of UA followed by lead citrate [30] for 5 min each.

### Acquisition, alignment, and analysis of serial tSEM images

Serial section images (8-bit TIFF; field size = 24,576 × 24,576 pixels) were acquired on a Zeiss Supra40 field emission scanning electron microscope in transmission mode (tSEM) [31] with ATLAS package, running at 28 kV, at 1.8 nm pixel size with 1.2 µs dwell time and 3.5 mm working distance. Serial tSEM images were aligned automatically using Fiji [32] (RRID:SCR_002285; http://fiji.sc) with the TrakEM2 plugin [33] (RRID:SCR_008954; http://www.ini.uzh.ch/~acardona/trakem2.html). The images were aligned rigidly first, followed by application of elastic alignment [34]. The aligned image stack was cropped to 14,424 × 19,512 pixels (image field size = 912 μm^2^) with Fiji/TrakEM2 and imported into Reconstruct [35] (RRID:SCR_002716; http://synapseweb.clm.utexas.edu/software-0) for 3D reconstruction and analyses. An image of grating replica (Electron Microscopy Sciences Cat# 80051), acquired along with serial section images, was used to calibrate pixel size. Mean section thickness was estimated based on the diameter of longitudinally sectioned mitochondria [36]. In serial tSEM images of sections that were immunolabeled, we first identified all axons containing any gold particles. An axon was considered as positively labeled if it contained gold particles (> 10 nm diameter after gold enhancement) outside mitochondria in at least two of three serial sections. To measure the density of enhanced gold particles, all particles were counted in four immunolabeled sections and categorized as positive labels or background. Particle densities were calculated from these counts divided by their respective areas in each of the analyzed sections. The density of axonal boutons was calculated based on the unbiased volume method [37], in which all boutons that did not intersect the exclusion planes were counted in a sub-volume of the 3DEM series encompassing 8 × 8 × 4.1 μm.

### Control for immunogold labeling

To control for non-specific binding of the primary and secondary antibodies, we collected serial thin sections from rAAV-injected tissue that was not labeled with tyramide-conjugated dye, but otherwise underwent immunogold labeling. This tissue was derived from the same slice as the one used for 3D reconstruction of a labeled axon as described above. In three consecutive sections (image field size = 1,520 μm^2^) from this control series, we counted axons containing at least one enhanced gold particle (> 10 nm diameter). None of these axons qualified as positively labeled, indicating the false positive rate is negligible.

An additional control was performed for non-specific antibody binding, in which another set of serial thin sections from rAAV-injected, tyramide-labeled tissue underwent immunogold labeling with the primary antibody omitted. These sections also showed no axons that qualified as positively labeled in the EM images. To control for self-nucleation of gold enhancement reagent, a tSEM image was also acquired from serial thin sections of the area CA1 that were not immunolabeled, but were incubated with the gold enhancement reagent as above and then stained with UA and lead citrate.

To measure the size of enhanced gold particles, the antibody conjugated with 5 nm gold was blotted on a PEI-coated TEM grid and then treated with the gold enhancement reagent as above. We acquired a tSEM image from a square field encompassing 4,096 pixels per side at 1.8 nm/pixel and thresholded the image to identify a total of 527 particles for their size measurement with the particle analysis function of Fiji. This measures the maximum caliper, which is the longest distance between any two points along the selection boundary. Enhanced particles that obviously formed from two or more 5 nm particles placed in close proximity, as evidenced by the presence of negative curvatures, were excluded from this analysis.

### Confocal image analysis

Fiji was used for processing and analysis of the confocal images. Maximum intensity projection images were generated from 5 optical sections encompassing 51.5 × 34.3 × 3.2 µm in x, y, and z. Image stacks from Cy5 channel were corrected for bleaching (the simple ratio method under bleach correction function in Fiji) before they were projected. Projected images from GFP and Cy5 channels were merged and thresholded to identify all puncta (≥ 100 nm diameter) labeled with either of the fluorophores, which were then assessed for co-expression in single channel images. A total of 682 labeled puncta were identified, excluding those at edges of the image. Acquisition of single channel image stacks for each of the fluorophores, rather than dual-channel stacks, caused a slight mismatch in their z-positions, which may have contributed to a small fraction of puncta to appear as single-labeled. The density of GFP-positive puncta was calculated from three additional confocal image stacks, encompassing 39.1 × 39.1 × 4.4 μm (two image stacks) or 58.6 × 58.6 × 3.2 μm (one image stack).

### Analysis of physiology recordings

The initial acquisition and analysis of EPSP were performed with SynchroBrain software (Lohmann Research Equipment). The initial maximum slope was measured over a 0.2-0.8 ms time frame that was held constant for all recordings in each slice. To calculate the magnitude of LTP, EPSP slopes were normalized to the average slopes obtained during the last 30 min of baseline recordings before the delivery of the first HFS. Then values across slices (mean ± SEM) were presented as times baseline. LTP magnitude at 30, 60, 120, and 180 min post-HFS was calculated by averaging the values for the preceding 20 min. Prism software package (Graphpad Software) was used for statistical analysis and plotting. The main tests performed were Student’s t-test and analysis of variance (ANOVA). Specific statistical tests and results are shown in the corresponding figure captions.

### Data availability

The following files generated and analyzed during the current study are deposited at Texas Data Repository (doi:10.18738/T8/QP43LB):

1. The original, unaligned serial tSEM images (VYLNH_raw1.zip [11.1 GB], VYLNH_raw2.zip [11.4 GB], and VYLNH_raw3.zip [12.0 GB])
2. The aligned 3DEM dataset (VYLNH_20181017.zip; 14.8 GB)
3. S1 Video (S1_video.mp4; 71.1 MB).

## Results

### Potentiation of synapses using light

We designed an adeno-associated virus vector (rAAV) for stimulating and tracking individual neurons and the trajectories of their axons. The construct encoded two proteins: a ChR2 [14] fused to GFP [15] and an APX from *P. sativum* [17] to generate the electron-dense deposits detected in EM. We chose ChR2^ET/TC^ based on its conductance and ability to sustain optical stimulation up to 60 Hz [14]. We modified APX to include amino acid substitutions for increased stability and enzymatic activity [5,6]. We enhanced its membrane and synapse targeting by adding a palmitoylation signal from growth-associated protein 43 (GAP-43) [18] to produce mAPEX2 (Fig 1A). A virus encoding ChR2 and mAPEX2 ensured that both proteins co-localized in the same cells. The observations from dissociated neurons and acute hippocampal slices reflect ChR2-GFP and mAPEX2 co-expression at the cellular and synaptic levels (Figs 1B and 1C).

**Fig 1.**
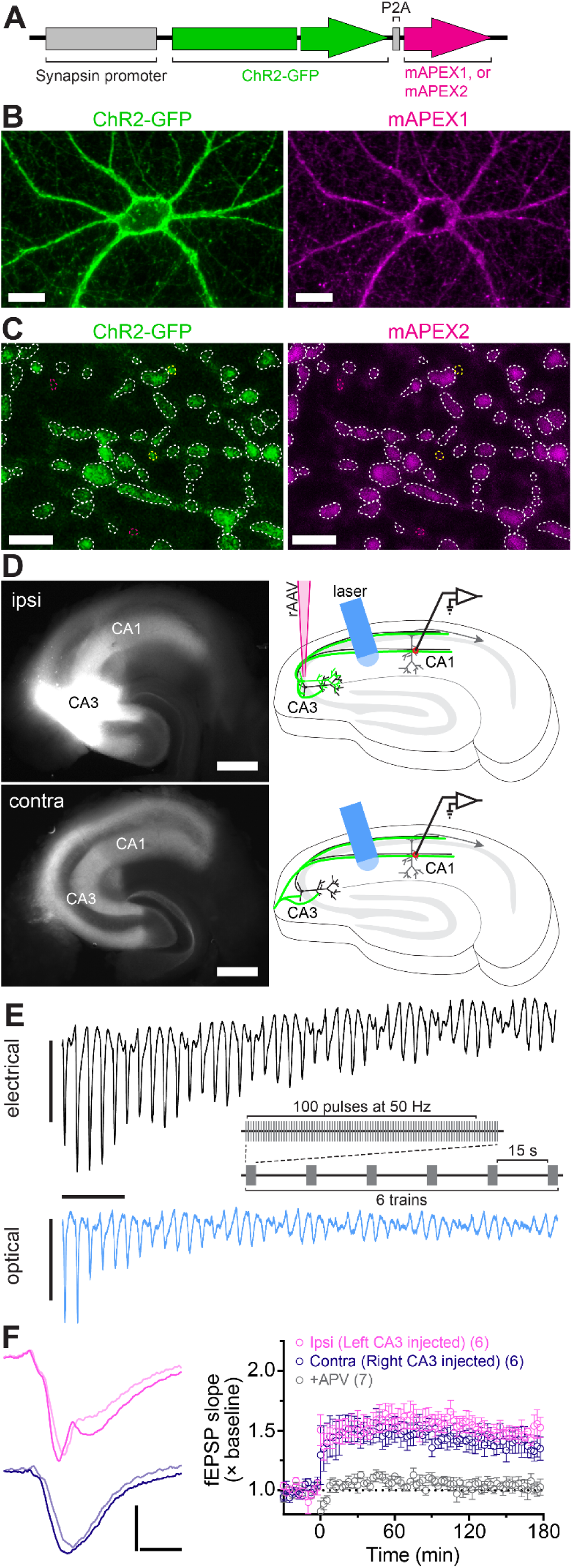
Viral expression of ChR2-GFP and mAPEX, and induction of optical L-LTP. (**A**) The rAAV was designed to co-express ChR2-GFP and mAPEX under a single synapsin promoter in neurons. During translation, the P2A peptide self-cleaves to yield the two proteins. (**B**) Cultured hippocampal neurons infected with the rAAV co-expressed ChR2-GFP and mAPEX1. Scale bars = 25 µm. (**C**) ChR2-GFP was co-expressed with mAPEX2 in the same axons in the area CA1 (dotted white lines). Yellow and magenta lines indicate ChR2-GFP only and mAPEX2 only puncta, respectively. Scale bars = 2 µm. (**D**) GFP fluorescence images (left) and experimental configurations (right) in ipsilateral (top, ipsi) and contralateral (bottom, contra) hippocampal slices. Scale bars = 500 µm. (**E**) Electrical (top) and optical (bottom) EPSPs evoked by a train of 50 Hz stimulations, recorded from CA1 middle *stratum radiatum* (C57B/6J strain). Responses to the first 40 pulses are shown and the stimulation artifacts are clipped from the eEPSP data. The inset shows HFS protocol for LTP induction. Scale bars = 1 ms, 2 mV (electrical), 1 mV (optical). (**F**) Optical HFS induced L-LTP in the Schaffer collateral and commissural pathways. *Left:* Representative oEPSP traces from ipsilateral (top) and contralateral (bottom) slices before (light shaded line) and 3 hr after (solid line) HFS. *Right:* Time course of oEPSP slope (mean ± SEM) showing optical L-LTP in ipsilateral (pink) and contralateral (purple) slices. Addition of APV blocked L-LTP (grey). The number of slices is indicated in parentheses. Scale bars = 4 ms, 1 mV.

We injected the rAAV vector into *stratum pyramidale* of the hippocampal area CA3 in one hemisphere of the adult mouse brain (S1 Fig, A). Epifluorescence microscopy confirmed robust expression of ChR2-GFP in the CA3 neurons and in Schaffer collaterals extending into ipsilateral area CA1. The contralateral hippocampus showed GFP-labeled commissural/associational fibers in the areas CA1 and CA3 (S1 Fig, B).

Four to six weeks after virus injection we prepared acute transverse slices from ipsilateral or contralateral hippocampus, four per hemisphere, covering the dorsal region. The slices were allowed to recover for 3 hr in interface chambers [38]. We then applied pulses of blue light (473 nm wavelength; ~14.5 mW power) via an optical fiber (300 μm diameter) positioned over the area CA1 *stratum radiatum* in each chamber (Fig 1D) and examined the ChR2 responses. Typically, 2-4 slices displayed optical responses. In these slices, optically-evoked field excitatory postsynaptic potentials (oEPSP) and population spike shapes recorded within different strata of area CA1 resembled the waveforms observed previously with electrical stimulation (eEPSP) [39–41], confirming a similar propagation of the signal. Although oEPSP were similar to eEPSP, we noticed a difference in the shape of the waveform: the initial activation stage consisted of more than one component, visible as change in slope. We assume this could be due to asynchrony of fiber firing at different sites along *stratum radiatum* because of the relatively larger area stimulated by light compared to concentric bipolar stimulating electrodes. In accordance with prior reports, oEPSP had comparable slopes and amplitudes between ipsilateral and contralateral groups of fibers [11,42], although these parameters were significantly smaller than in electrical responses (S1 Fig, D). Light-evoked short-term plasticity [43], measured as paired-pulse ratio (PPR), was also detected (S1 Fig, E). Optical responses were blocked by TTX (S1 Fig, F) and DNQX (S1 Fig, F), demonstrating their dependence on voltage-gated sodium channels and GluA receptors, respectively.

Next, we confirmed that ChR2 could follow trains of light pulses needed to induce L-LTP. Stimuli were delivered at 50 Hz (Fig 1E), a frequency sufficient for LTP induction [10,44] that also allows more time for ChR2 to recover between stimulation episodes [14] than the higher frequencies (100 Hz or higher, including theta-burst) typically used for electrical induction of LTP. Optical stimulation resulted in the trains of evoked oEPSP in CA1 *stratum radiatum* (Fig 1E, bottom) in a pattern comparable to the one evoked by 50 Hz electrical stimulation (Fig 1E, top). However, optical responses exhibited smaller initial amplitudes and underwent stronger desensitization during the trains of light stimuli.

Smaller amplitudes in our optical experiments could be due to incomplete ChR2 activation or to the presence of ChR2 in a relatively small subset of axons. To differentiate between these possibilities, we recorded oEPSP under varied light stimulus intensity. Input-output curves of oEPSP slope as a function of light intensity showed an apparent saturation at maximum level of ~14.5 mW (S2 Fig). Thus, our optical stimulation protocol maximized activation of all the targeted axons.

Optical stimulus regimes produced L-LTP that lasted for at least 3 hr (Fig 1F). Optically stimulated ipsilateral and contralateral slices showed the same degree of potentiation. However, the likelihood of achieving L-LTP varied, and the success rate was 67% and 42% for ipsilateral and contralateral slices, respectively. Optical L-LTP induction was blocked by application of 2-amino-5-phosphonovaleric acid (APV), reflecting a dependence on GluN receptors (Fig 1F).

To isolate the rAAV-targeted CA3 axons as the sole source of excitatory synaptic transmission and to avoid possible recruitment of unlabeled fibers, we recorded oEPSP from contralateral slices with the area CA3 severed pre-recovery (Fig 2A). Using contralateral slices additionally eliminated the likelihood that CA1 neurons could be labeled with ChR2 and stimulated independently of the CA3 axons. Severing CA3 had little effect on the likelihood or magnitude of L-LTP from electrical stimulation (Fig 2B). In contrast, the induction rate was reduced from 42% to 33% in contralateral sections with severed CA3, and the magnitude of LTP was smaller at 1 hr compared to intact sections, but nearly identical by 3 hr (Figs 2C and 2D). The magnitude of electrical LTP in intact and cut slices was similar at 1 hr and 3 hr post-stimulation (Figs 2B and 2D). We conclude that the optical stimulation of genetically specified CA3 commissural fibers is sufficient to induce L-LTP in the area CA1.

**Fig 2.**
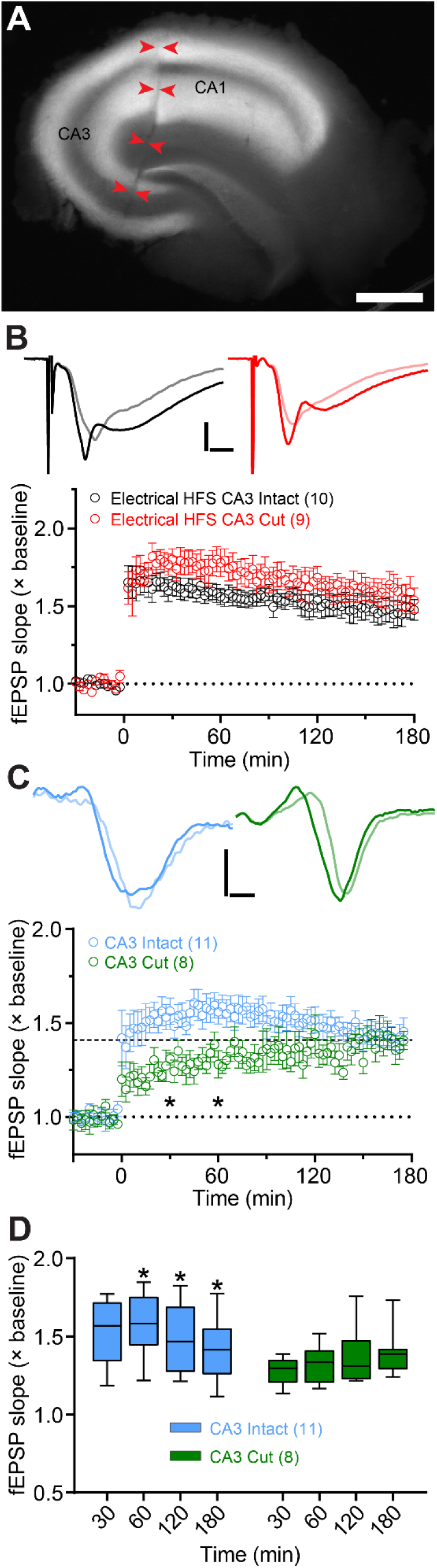
Optical stimulation of CA3→CA1 commissural fibers was sufficient for L-LTP. (**A**) Representative image of a contralateral slice with CA3 cut off from CA1. Scale bar = 500 µm. (**B**-**C**). LTP induced by electrical (**B**) and optical (**C**) HFS in intact and cut slices. The example traces show EPSPs recorded before (light shaded lines) and 3 hr after HFS (solid lines). Optical LTP magnitude (**C**) at 30 and 60 min post-HFS was significantly different between intact and cut slices (oEPSP slope F_(1, 17)_ = 4.74, p < 0.05; Time F_(3, 51)_ = 0.95, p > 0.05; Interaction F_(3, 51)_ = 6.60, p < 0.0001; repeated measures two-way ANOVA with Bonferroni’s post-hoc tests). Scale bars = 2 ms, 2 mV (**B**); 1 ms, 0.5 mV (**C**). (**D**) Summary data for oEPSP slope change at different time points following optical HFS in intact and cut slices. The intact slices showed significant changes in LTP magnitude in the last 120 min of recordings (F_(1.28,12.8)_ = 5.75, p < 0.05; repeated measures one-way ANOVA). The box plots show medians and interquartile ranges, with whiskers extending from minimum to maximum values. The number of slices used for each condition is indicated in parentheses.

### *Post hoc* labeling and reconstruction of potentiated synapses

Post hoc 3DEM analysis of activated synapses and subcellular organelles requires well-preserved slice tissue, which is typically achieved by chemical fixation with glutaraldehyde. Thus, the expressed EM tag must retain its enzymatic activity to produce electron-dense deposits in fixed samples. Like the original APEX [5,6], we verified that mAPEX2 was active after glutaraldehyde fixation by observing conversion of 3,3’-diaminobenzidine (DAB) into osmiophilic polymers, which appear dark brown under light microscopy or as amorphous electron-dense deposits under EM (S3 Fig).

We fixed brain slices that displayed L-LTP following optical stimulation at the 3 hr time point and had robust GFP labeling (Fig 3A, Step 1; see Materials and Methods). Orthogonal vibratome sections spanning area CA1 (Fig 3A, Steps 2-3) were incubated with tyramide conjugated with Alexa Fluor 647. Membrane-associated mAPEX2 then catalyzed the tyramide signal amplification reaction (TSA) locally upon the addition of H_2_O_2_ (Fig 3A, Step 4). After heavy metal staining and epoxy embedding, serial thin sections (~60 nm thickness) were collected from a region of interest containing the middle of *stratum radiatum* (Fig 3A, Steps 5-7). For the series shown in Figs 3A (Step 7), 3C, and 4, a total of 72 sections were collected from portions of tissue at least 10 µm from the surface of a 50 µm thick vibratome section.

**Fig 3.**
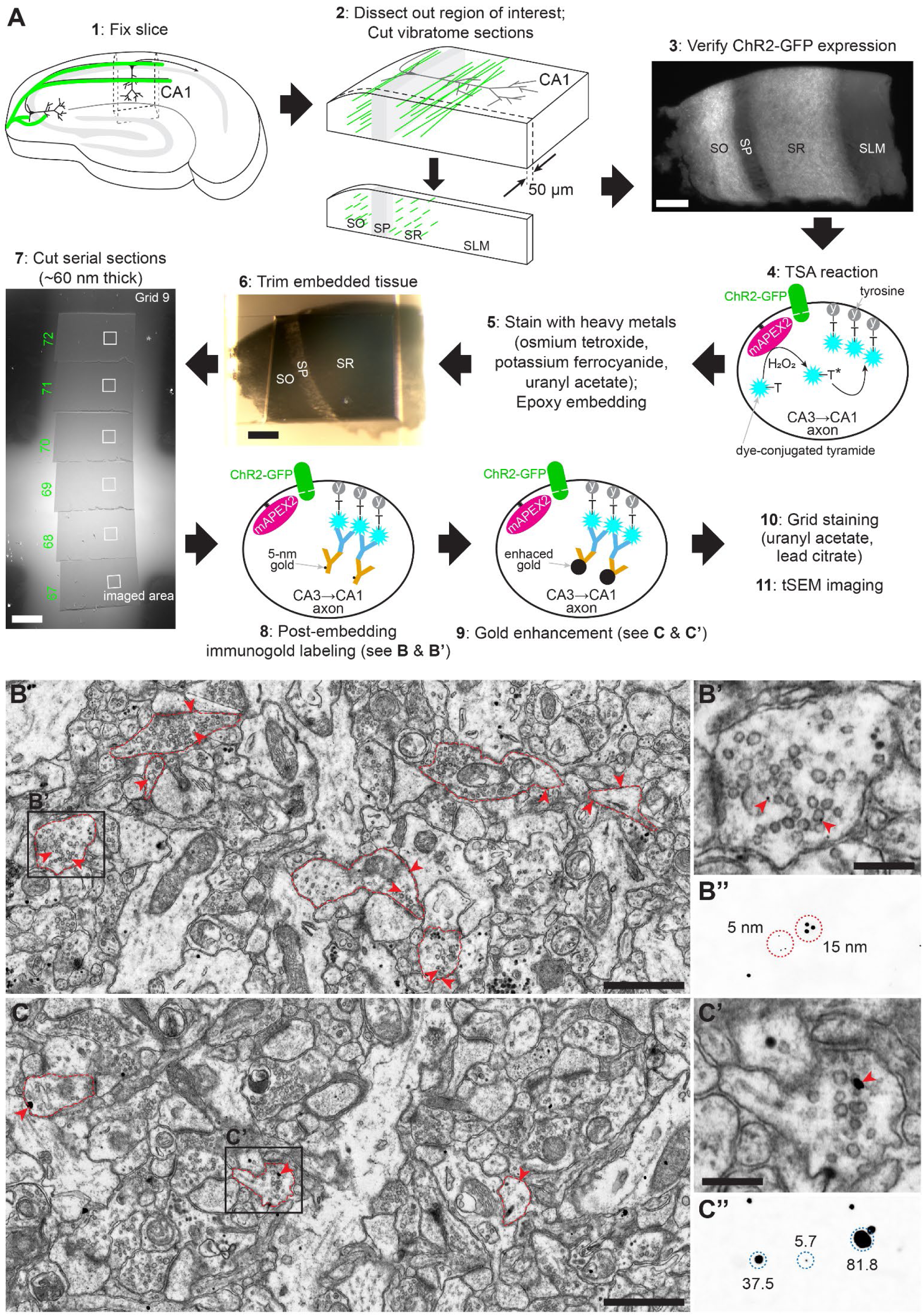
mAPEX2-catalyzed labeling and 3DEM identification of rAAV-targeted axons. (**A**) Workflow for processing of hippocampal slices following optical L-LTP. Vibratome sections from the fixed area CA1 underwent tyramide signal amplification (TSA) catalyzed by mAPEX2 to deposit Alexa Fluor dye in the targeted axons. The dye-labeled section was then stained with heavy metals and embedded into epoxy before being cut into serial thin sections. The dye-containing axons in a subset of the sections were immunolabeled with 5 nm gold particles, followed by gold enhancement. Scale bars = 100 µm. (**B** and **B’**) A low magnification tSEM image of the area CA1 *stratum radiatum* after immunolabeling (Step 8 in **A**). Axonal boutons indicated by red contours were positively labeled with 5 nm gold particles (red arrowheads). Area indicated by black rectangle is enlarged in **B’**. Scale bar = 1 µm in **B**, 250 nm in **B’**. (**B’’**) A tSEM image of colloidal gold particles (5 nm and 15 nm). To visualize the 5 nm particles more clearly, this image was acquired at 1 nm/pixel and scaled to the same magnification as **B’**. (**C** and **C’**) Same as **B** and **B’**, imaged after gold enhancement (Step 9 in **A**). Scale bar = 1 µm in **C**, 250 nm in **C’**. (**C’’**) A tSEM image of enhanced gold particles. The numbers indicate diameters in nm (also see S5 Fig 5).

In a subset of sections, at the beginning and end of the series, we labeled the axons expressing mAPEX2 that now contained the dye-tyramide molecules with anti-dye antibodies, then gold-conjugated secondary antibodies (Fig 3A, Step 8). This limited labeling ensured that the ultrastructure was visible while having sufficient labeling to identify the genetically-targeted axons unambiguously (Figs 3B and 3B’). We considered an axon to be positively labeled if it contained gold particles in at least two of three serial sections per grid. We tested 5 nm and 15 nm colloidal gold particles. Staining with smaller particles suffered from relatively low signal-to-noise, limited visibility of the gold particles (nominal particle size = 5 nm vs. pixel size = 1.8-2.0 nm), and difficulty in distinguishing the particles from artifacts and subcellular features of similar size (Fig 3B’ and S4 Fig). Working with 15 nm gold particles improved particle visibility (Fig 3B’’) but reduced labeling sensitivity, making labeled axons harder to identify. We boosted the visibility of 5 nm particles by increasing their size with a gold enhancement technique [45] (Fig 3A, Step 9). The enhanced particles were of high contrast and uniformly electron-dense with smooth edges, which made them distinct from artifacts and subcellular structures of similar size (e.g., glycogen granules, darkly stained membrane, and precipitation from post-section staining; Figs 3C, 3C’, and S4 Fig). Enhancement of immunogold particles blotted on a blank grid resulted in particles with diameter ranging from 2.5 nm to 85.3 nm (median = 37.1 nm), significantly improving their ease of identification (Fig 3C’’ and S5 Fig). Control sections devoid of immunogold labeling, but treated with the enhancement reagent, showed electron-dense particles with the diameter ≤ 10 nm, likely resulting from reagent self-nucleation (S5 Fig). We therefore excluded these small particles during identification of labeled axons. Thus, in the gold enhanced material, our criterion for positively labeled axons was the presence of particles > 10 nm in diameter in at least two of three consecutive sections.

In this 3DEM series, 23 axons (2 axons per 100 μm_2_) were positively labeled, and 170 axons (19 axons per 100 μm_2_) contained enhanced gold particles but did not meet the criteria. The 23 axons that were confirmed positive could include true and false positives, while the 170 negative axons would reflect true and false negatives. To estimate the numbers in each category, we performed control immunogold labeling on serial thin sections from the same rAAV-injected tissue that otherwise did not undergo labeling with the tyramide-conjugated dye. In these sections, we found 223 axons containing at least one enhanced gold particle (15 axons per 100 μm_2_). None of these axons qualified as positively labeled, indicating that the false positive rate is negligible. Thus, we deemed as true negatives the 15 of 19 axons per 100 μm_2_ that did not meet the criteria in the tyramide-labeled series. This suggests that the remaining 4 of 19 axons could be false negatives under our current labeling protocol for 3DEM, and that positively labeled axons could represent ~30% (2 positively labeled out of 6 containing gold particles) of rAAV targeted axons.

We also measured the density of boutons belonging to positively labeled axons in the 3DEM series (12 per 1,000 μm^3^), and that of GFP-positive puncta in three confocal image volumes (mean = 35 per 1,000 μm^3^; range = 20 to 47 per 1,000 μm^3^). Although measured from tissue samples from different animals, these observations also suggest that the confirmed positives could represent ~30% of the rAAV targeted axons. However, frequency of potential false negatives relative to the total number of axonal boutons is small (~1%, or about 10-30 of 2604 boutons per 1,000 μm^3^). It will be possible to uncover ultrastructural correlates of potentiation by comparing labeled axons from slices that have and have not received optical stimulations. To show that the plasticity-related ultrastructural changes are restricted to the positively labeled axons, one could also analyze unlabeled axons from slices with or without optical stimulations. In this case, the contribution of false negatives to the pool of unlabeled axons in either preparation will be extremely low, on the order of 1%.

The density of positive particles within labeled axons after gold enhancement was 2.85 ± 0.88 per µm^2^, while the overall background particle density was 0.33 ± 0.02 per µm^2^ (mean ± SEM; n = 4 sections). Thus, the signal-to noise ratio was 8.7.

Finally, we reconstructed a labeled axon from an rAAV-infected CA3 neuron forming synapses onto dendritic spines in middle *stratum radiatum* of the area CA1 from a slice that showed optically induced L-LTP (Fig 4 and S1 Video). This axon contained gold particles in the first three sections and three of the last four sections that were immunolabeled (Figs 4A and 4C). This axon had two multi-synaptic boutons, one of which was reconstructed only partially because it was at one edge of the serial section series (Fig 4B). This partial bouton formed synapses with two spines (s1 and s2 in Fig 4B; also see S1 Video). One of these spines (s1) had another branch that formed a synapse with a separate axonal bouton that was unlabeled. The synapses on the branched spine (s1) were both macular and of similar sizes (0.029 µm^2^ and 0.021 µm^2^). The bouton that was completely reconstructed contained two mitochondria and formed synapses with two spines from different dendrites (Fig 4B; S1 Video). One of these synapses was perforated (area = 0.11 µm^2^), while the other was macular (area = 0.037 µm^2^). These findings are consistent with synapses on this axon having been potentiated [46].

**Fig 4.**
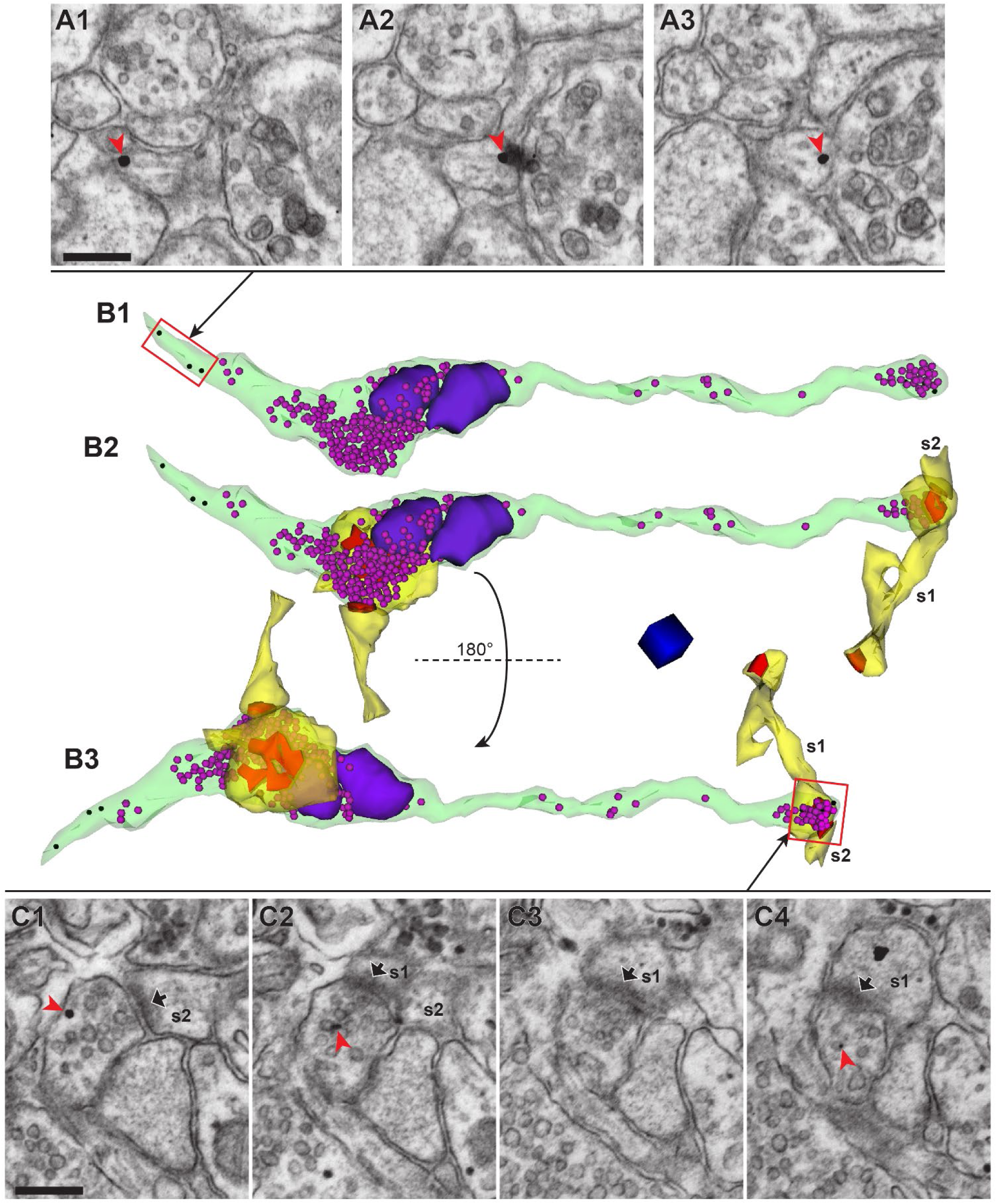
Post-embedding immunogold labeling for Alexa Fluor dyes deposited by mAPEX2-driven TSA reaction allowed for 3DEM identification of rAAV-targeted axons, while maintaining excellent ultrastructure. (**A1**-**A3**) Three adjacent serial tSEM images of an axon containing immunogold labeling (red arrowhead). Scale bar = 250 nm. (**B1**) 3D reconstruction of the labeled axon (green) shown in **A**, which contained immunogold labels (black spheres) and synaptic vesicles (magenta spheres). Two mitochondria (purple) were associated with one of the two boutons. Red rectangle represents a portion of this axon shown in **A1**-**A3**. (**B2**) Same axon as **B1**, with reconstructions of spines (yellow) forming synapses (red) with this axon. Note, s1 was a branched spine with one of the heads forming a synapse with another axon. The second spine (s2) and its PSD could be reconstructed only partially because they were located at the end of the tSEM image series. Both axonal boutons are multi-synaptic, with each bouton forming synapses with two spines originating from different dendrites. (**B3**) Same as **B2**, rotated along the horizontal axis 180° to provide a different view of synapses and spines. One of the synapses at mitochondria-containing bouton was perforated (also see S1 Video). Red rectangle represents a portion of this axonal bouton shown in **C1**-**C4**. Scale cube = 250 nm per side. (**C1**-**C4**) Four adjacent serial tSEM images of the gold-labeled (red arrowhead) axonal bouton, forming synapses with two dendritic spines (s1 and s2; PSDs indicated by black arrows). Scale bar = 250 nm.

## Discussion

Genetic targeting of specific neuron populations for imaging and activation has transformed structure-function studies of brain circuitry. Until now, however, tools for examining the ultrastructure of genetically specified neurons post-manipulation have been lacking. Here we describe methods for inducing activity-dependent plasticity in a genetically defined subset of neuronal synapses and for identifying optogenetically potentiated axons. We demonstrate light-evoked L-LTP in acute hippocampal slices. Previous ultrastructural studies of potentiated synapses examined effects across differentially activated populations of synapses, but the histories of individual synapses were unknown. The unilateral infection of CA3 neurons yields CA3→CA1 projections that co-express a light-dependent actuator and an EM tag from a single rAAV amid a larger population of unlabeled axons. Hence the advance of our approach is that recently potentiated synapses can be compared to neighboring unstimulated synapses in the same block of tissue.

### Light stimulation of CA3 axons resulted in robust oEPSP in ChR2-expressing hippocampal slices

In hippocampal slices prepared from rAAV injected animals, light pulses induced oEPSP that depended on voltage-gated sodium channels and GluA receptors. The light intensity used for optical high-frequency stimulation (HFS) was shown to achieve the maximal responses. Furthermore, the expressed ChR2 reliably responded to light trains delivered at 50 Hz, a frequency sufficient to produce L-LTP [10,44]. Together these findings ensure that synapses identified by the *post hoc* 3DEM analysis were activated during the optogenetic LTP induction protocol.

### Optical L-LTP was produced in intact hippocampal slices expressing ChR2-GFP and mAPEX2

We used high-frequency optical stimulation to induce GluN receptor-dependent L-LTP lasting at least 3 hr in a subpopulation of CA3→CA1 synapses containing presynaptically expressed ChR2. Recent reports suggest molecular and synaptic asymmetry between right and left hippocampus affecting LTP induction and endurance [11,47]. These findings prompted us to explore induction and expression of optical L-LTP in both hemispheres. The rAAV was injected into the left (ipsilateral) or right (contralateral) hippocampus, and all slices were prepared from the left hemisphere. L-LTP reached similar magnitudes whether the ipsilateral Schaffer collaterals or contralateral commissural axons were optically stimulated.

This discrepancy with the prior papers could be due to technical differences, including recording configurations (field vs. single-cell recordings) or plane of hippocampal slices (transverse vs. coronal). Our findings are also consistent with prior studies using electrical stimulation. For example, in rat hippocampal slices where Schaffer collaterals or commissural fibers were isolated by kainic acid lesion, the properties of LTP produced independently by each pathway were similar to those observed in unlesioned slices [42,49]. Recent *in vivo* studies with intact brain also detected a robust LTP mediated by Schaffer or commissural axons [49].

### Optical L-LTP did not require recruitment of unlabeled axons

In order to make direct comparisons between synapses with different activation histories, optical HFS must activate only the rAAV-targeted axons. In the intact contralateral system, the measured oEPSP responses could be contaminated if back-propagating action potentials from labeled commissural fibers were to recruit neighboring unlabeled CA3 neurons through recurrent collaterals. To test whether our oEPSP responses had been amplified through light-independent intra-hippocampal connections, we compared optical stimulation in contralateral slices with intact or severed area CA3. Compared to intact slices, the cut slices had a lower success rate for L-LTP (42% vs. 33%), yet these results suggest L-LTP could certainly be achieved without recruitment of unlabeled axons.

There was a slower rise to maximum LTP magnitude in the cut slices that might have been caused by reduced synchronicity of firing if unlabeled axons are recruited in the intact system. Other experiments have witnessed this slow rise to LTP. For example, LTP can be produced with a slow onset with frequencies normally used for induction of LTD (1-5 Hz) [50]. Sometimes weak presynaptic stimulation paired with depolarization of postsynaptic neurons results in LTP with a similar slow rise early during expression [51,52]. Induction of mGluR-dependent [53] or BDNF-dependent LTP [54] both show slow rises in LTP magnitude. Furthermore, at the developmental onset of L-LTP the expression can also have a slow rise [38]. Each of these conditions is also submaximal in the induction paradigms, not unlike activation of less than all the axons with light versus electrical stimulation. However, since both intact and cut slices expressed optically induced L-LTP of the same magnitude at 2 and 3 hr post induction, the recruitment of unlabeled axons was unlikely to be the critical factor.

### mAPEX2 labeled axons are compatible with ultrastructural analysis of synapses

To study synaptic connectivity and function, it is necessary to know which synapses were engaged. This level of analysis has been limited by the availability of ultrastructural tools to identify activated synapses. We aimed to label the targeted axons discretely without obscuring or compromising the integrity of their subcellular and synaptic components. Since the genetically encoded mAPEX2 was expressed in the tissue and compatible with microwave-enhanced chemical fixation containing glutaraldehyde, it replaced the use of pre-embedding antibody labeling to identify rAAV-targeted axons. mAPEX2 accommodates tyramide-conjugated fluorescent dyes, which are then deposited in the targeted cells by the TSA reaction [55]. The combination of mAPEX2-driven TSA reaction and post-embedding immunogold labeling allowed reliable identification of the targeted cells throughout the 50 µm vibraslices, at depths not accessible by pre-embedding antibody methods (< 10 µm). These features revealed optimal preservation of ultrastructure in acute slice tissue for analyses of synapses and organelles after targeted optogenetic manipulations. We show that this process is compatible with conventional fixation, processing, epoxy infiltration, and post-embedding immunogold labeling [56,57]. We show that only a subset of serial sections need be labeled to track axons from the targeted cells, and even in the labeled sections, the particles did not obscure objects of interest.

L-LTP induced by electrical theta-burst stimulation in the area CA1 is associated with an increase in the abundance of multi-synaptic boutons in the population of stimulated axons [46]. The 3D reconstruction of a labeled axon from an optogenetically potentiated slice showed two neighboring multi-synaptic boutons, consistent with this prior report, but now from an identified axon. Future work will address whether such shifts in synapse configurations are dependent on activation history.

The methods described here provide reliable physiology and labeling compatible with conventional tissue fixation and the processing techniques needed to preserve subcellular organelles. They make an ideal approach to link synapse ultrastructure and function in intact circuits of genetically defined neurons.

## Acknowledgements

The authors wish to thank the following colleagues for their help with this work: Stefanie Esmond and Bridget Kajs for rAAV preparation; Geoff Dilly, Melissa Burks, Molly O’Gara for stereotaxic rAAV injections; John Mendenhall for help with EM and useful discussions on labeling strategies; Dan Johnston for use of a Zeiss light microscope and helpful discussions on electrophysiology; Nuno de Costa (Allen Institute for Brain Science) for Ni-DAB protocol; Jung-Hwa Tao-Cheng (NINDS Electron Microscopy Facility) for suggesting the use gold enhancement; Patrick Parker for help in preparing this manuscript. All are at the University of Texas at Austin unless otherwise noted. The authors also acknowledge the following funding sources: Brain Research Foundation Scientific Innovations Award (to K.M.H.), NSF Grant (1707356 to K.M.H.), NIH Grants (MH095980, R56MH095980, MH104319 and NS074644 to K.M.H.; EY026446, EY026442, and NS094330 to B.V.Z.), Human Frontier Science Program (RGP0041 to B.V.Z.), and The University of Texas System UT BRAIN Seed Grants (NNRI 365222 and 365289 to B.V.Z.).

## Supporting information

**S1 Fig. Verification of rAAV injections and further characterization of oEPSP at CA3→CA1 synapses.** (**A**-**B**) Unilateral injections of rAAV into the area CA3 result in GFP labeling of Schaffer collaterals and commissural fibers. (**A**) A diagram (modified from ref. 58) showing the approximate site of rAAV injection site in the mouse hippocampal area CA3. An infected CA3 neuron (green) on the right hemisphere depicted under the injection needle (magenta) projects its axons to synapse onto neurons in both ipsilateral and contralateral CA1 via Schaffer collaterals and commissural fibers, respectively. These axons can also synapse onto uninfected CA3 neurons (black). (**B**) A montage of two epifluorescence images of a coronal section through the injection site in the right area CA3. The right hemisphere (injected side) was partially damaged during extraction of the brain. Scale bar = 1 mm. (**C**-**F**) In acute transverse slices of the hippocampus prepared from rAAV injected mice, light pulse stimulation of the virally targeted CA3 axons evoked oEPSPs in the area CA1. (**C**) Example traces of oEPSPs recorded from an electrode positioned in different CA1 layers (colored circles): strata oriens (SO), pyramidale (SP), radiatum (SR), and lacunosum-moleculare (SLM). Optical fiber was placed in SR in the proximal area CA1, ~400 µm from the recording electrode. Scale bars = 5 ms, 1 mV. (**D**) Amplitude (left) and slope (right) of optically and electrically evoked field EPSPs. Optical stimulation was delivered at the maximum light intensity, while electrical stimulation was at half-maximum. Horizontal lines and error bars indicate mean ± SEM. The number of slices used for each condition is indicated in parentheses. (**E**) Optical paired-pulse stimulation induces slight facilitation (mean ± SEM, n = 6 slices). Scale bars = 10 ms, 1 mV. (**F**) oEPSP was blocked by application of 1 µM TTX (left) or by 20 µM DNQX (right). Scale bars = 5 ms, 1 mV (TTX); 5 ms, 2 mV (NBQX). The recordings shown in **D**-**F** were made from the middle SR.

**S2 Fig. Input-output (IO) curves recorded from the area CA1 in intact and cut slices.** (**A**) Representative oEPSP traces recorded from a slice with intact CA3. (**B**) Same as **A**, but recorded from a cut slice. Scale bars = 1 ms, 1 mV for **A** and **B**. (**C**) Optical IO curves recorded from intact and cut slices. The intensities of light stimulation used to record optical IO: 30% (~4 mW), 60% (~9mW), 90% (~13mW), and 100% (~14.5 mW). There is no significant difference between responses evoked by 90% and 100% light intensities (t = 0.317, df = 10, p > 0.05 for intact slices; t = 0.342, df = 9, p > 0.05 for cut slices; two-sided paired t-test). (**D**) Summary of oEPSP slope (left) and amplitude (right) recorded at the maximum light power. (**E**) Input-output curves recorded using an electrical stimulation. (**F**) Summary of eEPSP slope (left) and amplitude (right) recorded at intensity that resulted in approximately half-maximum slope. All graphs (**C**-**F**) show mean ± SEM. Different sets of slices were analyzed for **C** and **D**; **E** and **F**. The number of slices used for each condition is indicated in parentheses.

**S3 Fig. Enzymatic activity of mAPEX is preserved after chemical fixation with glutaraldehyde.** Diaminobenzidine (DAB) was used as a substrate because autofluorescence from glutaraldehyde makes it difficult to assess labeling with tyramide-conjugated fluorescent dyes. (**A**) Right: mAPEX1 expressed in dissociated rat hippocampal neurons, fixed with 6% glutaraldehyde and 2% formaldehyde, were capable of generating the dark brown DAB reaction product. Left: Control neurons fixed and treated with DAB in the same manner did not exhibit the reaction product. Scale bars = 50 µm. (**B**) Two serial tSEM images showing axons labeled with Ni-enhanced DAB (red contours) through the area CA1 from a perfusion-fixed C57B/6J mouse. The rAAV was injected into the ipsilateral hippocampal area CA1 to express mAPEX2. The fixative contained 2.5% glutaraldehyde and 2% formaldehyde. Scale bars = 500 nm. Insets: Enlarged areas indicated by black rectangles. Electron-dense Ni-DAB reaction product obscures subcellular structures in the labeled axons. In contrast, small synaptic vesicles are visible in an unlabeled axonal bouton nearby (ax). Scale bars = 100 nm.

**S4 Fig. Electron-dense artifacts and subcellular structures present in unlabeled sections.** Electron-dense artifacts and subcellular structures present in unlabeled sections. Five serial tSEM images from the series presented in Fig 3C. These images were acquired from serial thin sections that were not immunolabeled for Alexa Fluor dye, but stained with uranyl acetate and lead citrate (UA/Pb) prior to tSEM imaging. While glycogen granules fill glial processes (green arrowheads in ‘g’), they are less common in axons (ax) and boutons (b1, b2, and b3) and do not appear in the same profiles over two consecutive sections (yellow arrowheads). Post-section staining with UA/Pb can be a source of electron-dense artifacts of various size and shape (purple arrowheads) that are usually confined to single sections. On rare occasions, small artifacts could appear in two consecutive sections. However, their small size (diameter ≤ 10 nm) is similar to those generated as a result of background nucleation during gold enhancement (see S5 Fig), and therefore, can easily be distinguished from positive labeling should they also occur in immunolabeled sections. Scale bar = 250 nm.

**S5 Fig. Electron-dense artifacts due to gold enhancement reagent. Electron-dense artifacts due to gold enhancement reagent.** (**A**) A histogram showing the distribution of diameters of gold-enhanced gold particles (bin size = 2 nm). A total of 527 particles were analyzed with Fiji software (see Materials and methods). Their diameter ranged from 2.5 nm to 85.3 nm with the median of 37.1 nm and the mean of 36.5 ± 0.61 nm (SEM). A vertical line at 10 nm indicates the cutoff for gold particles excluded during identification of labeled axons. (**B**) A tSEM image acquired from serial thin sections that were not immunolabeled, but incubated with the gold enhancement reagent and then stained with uranyl acetate and lead citrate. Small electron-dense particles (indicated by dotted red circles) were 10 nm or smaller in diameter, and were present in dendrites (den), axons (ax), and boutons (b). These small particles likely formed as a result of self-nucleation of gold enhancement reagent, and therefore were excluded during identification of labeled axons. Scale bar = 250 nm.

**S1 Video. A video of the reconstructed axon shown in** Fig 4. File name: S1_video.mp4. File size: 71.1 MB. This video is also available as an external file – see Data Availability.

