## Supplementary figures and images for "Ultrastructure of light-activated axons following optogenetic stimulation to produce late-phase long-term potentiation"

### Supplemental Figure 1

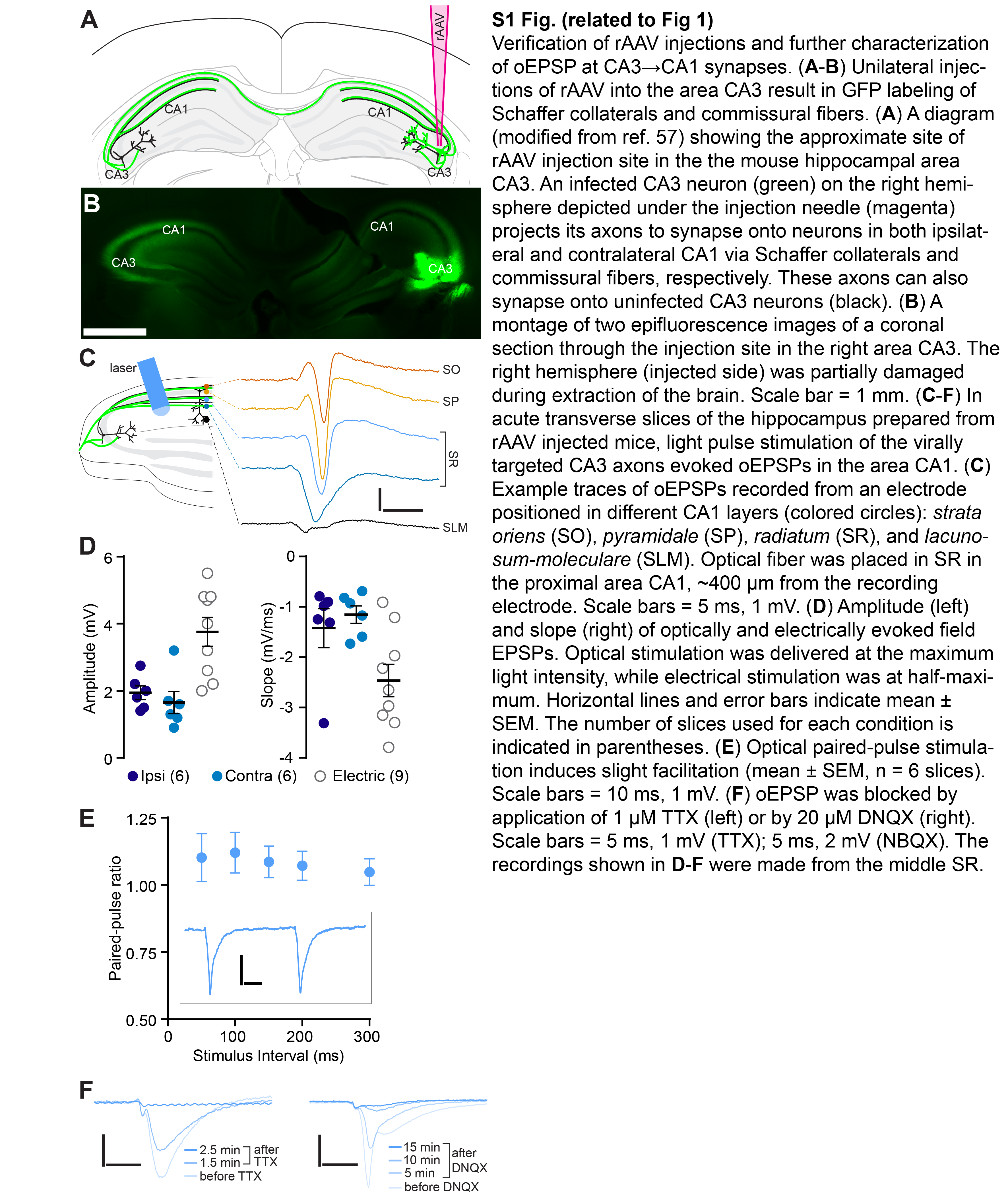

### Supplemental Figure 2

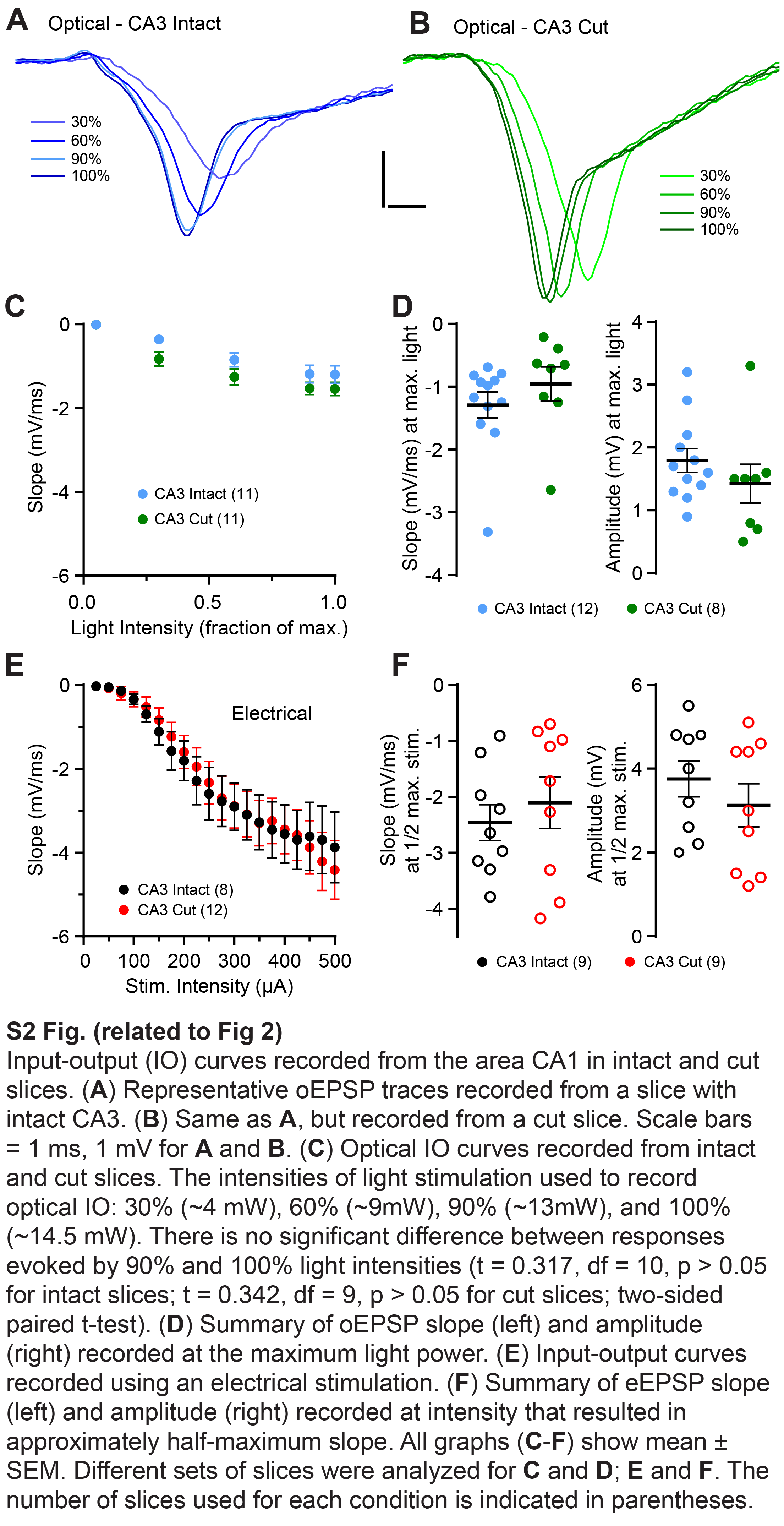

### Supplemental Figure 3

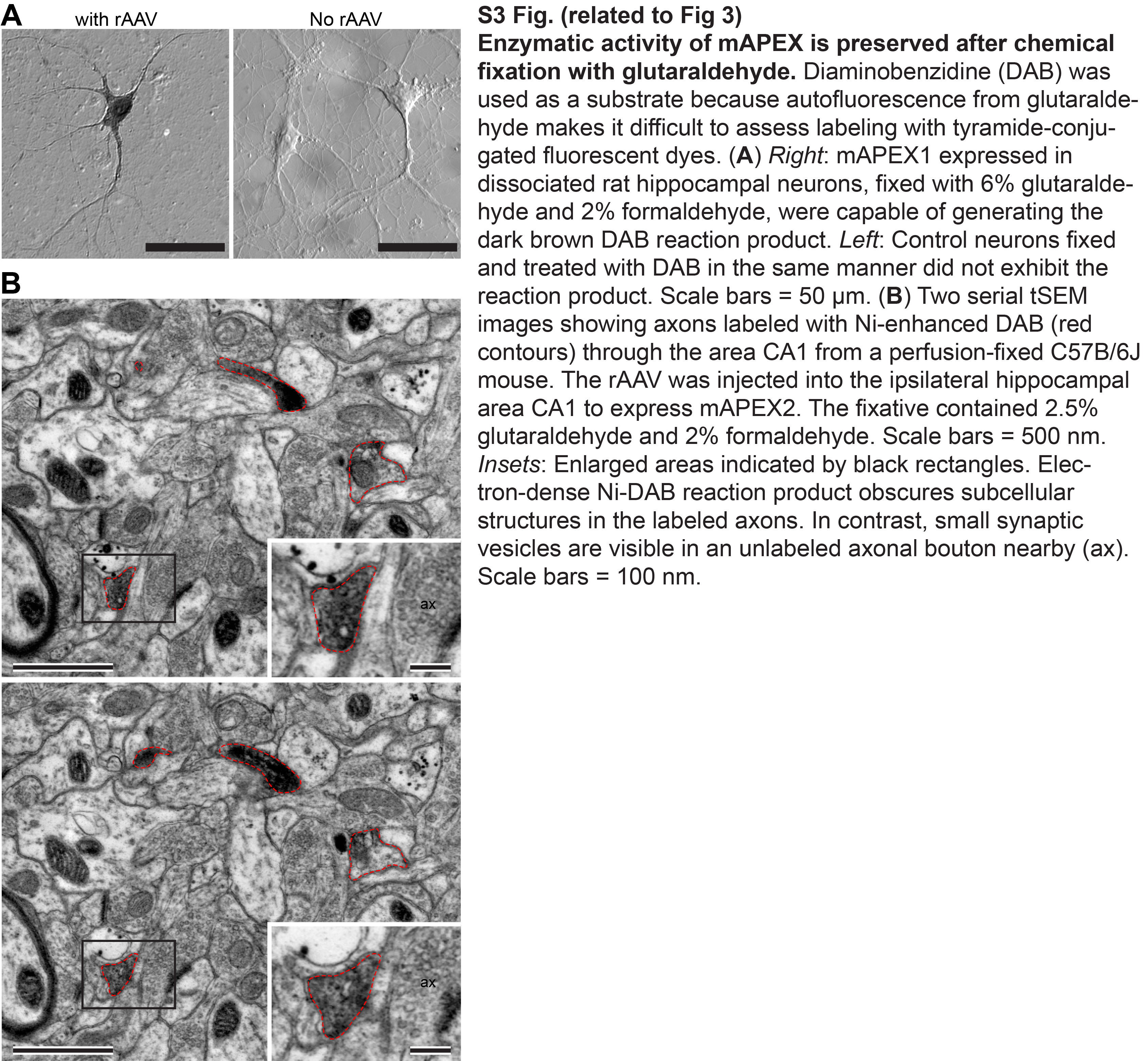

### Supplemental Figure 4

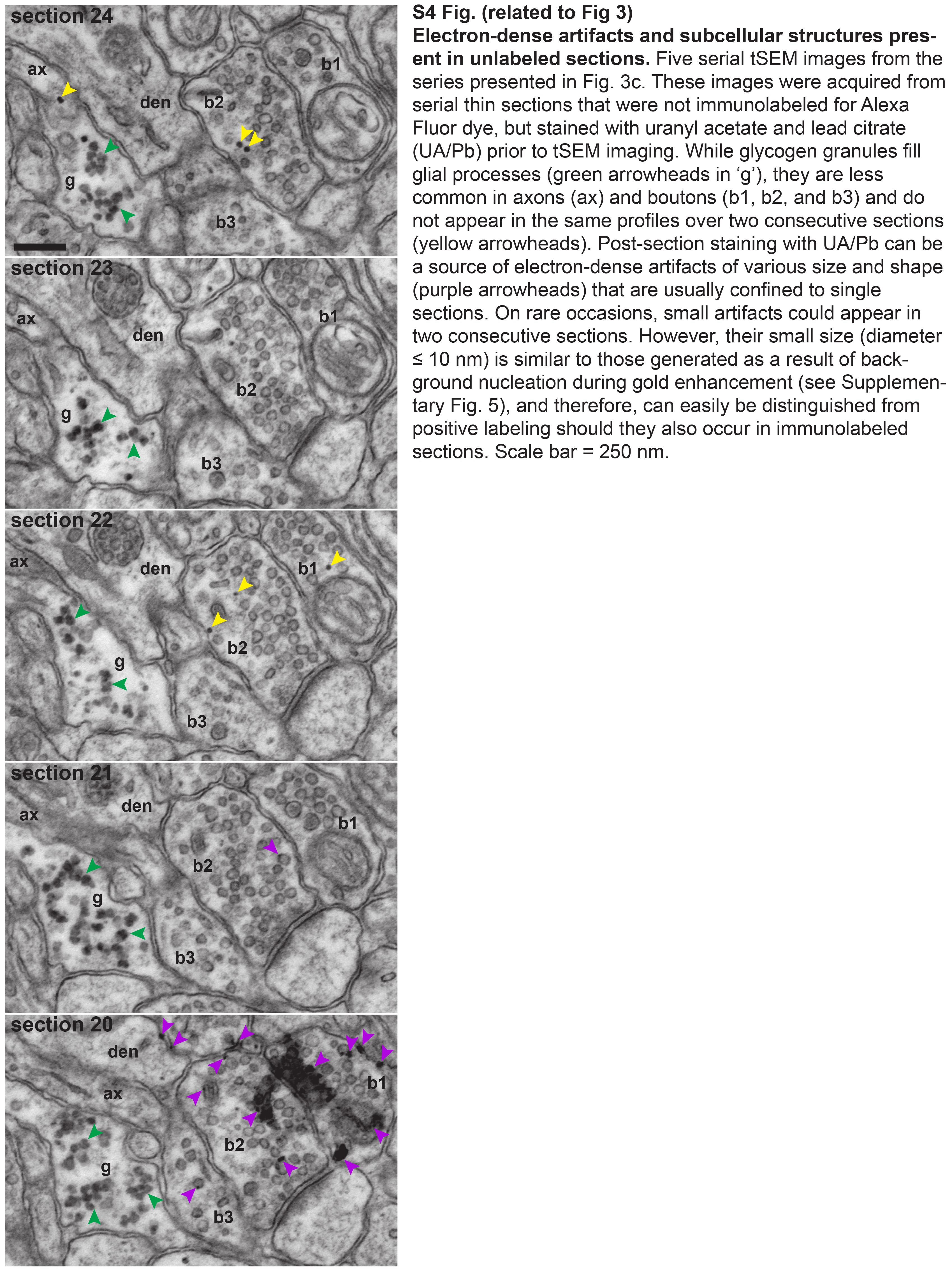

### Supplemental Figure 5

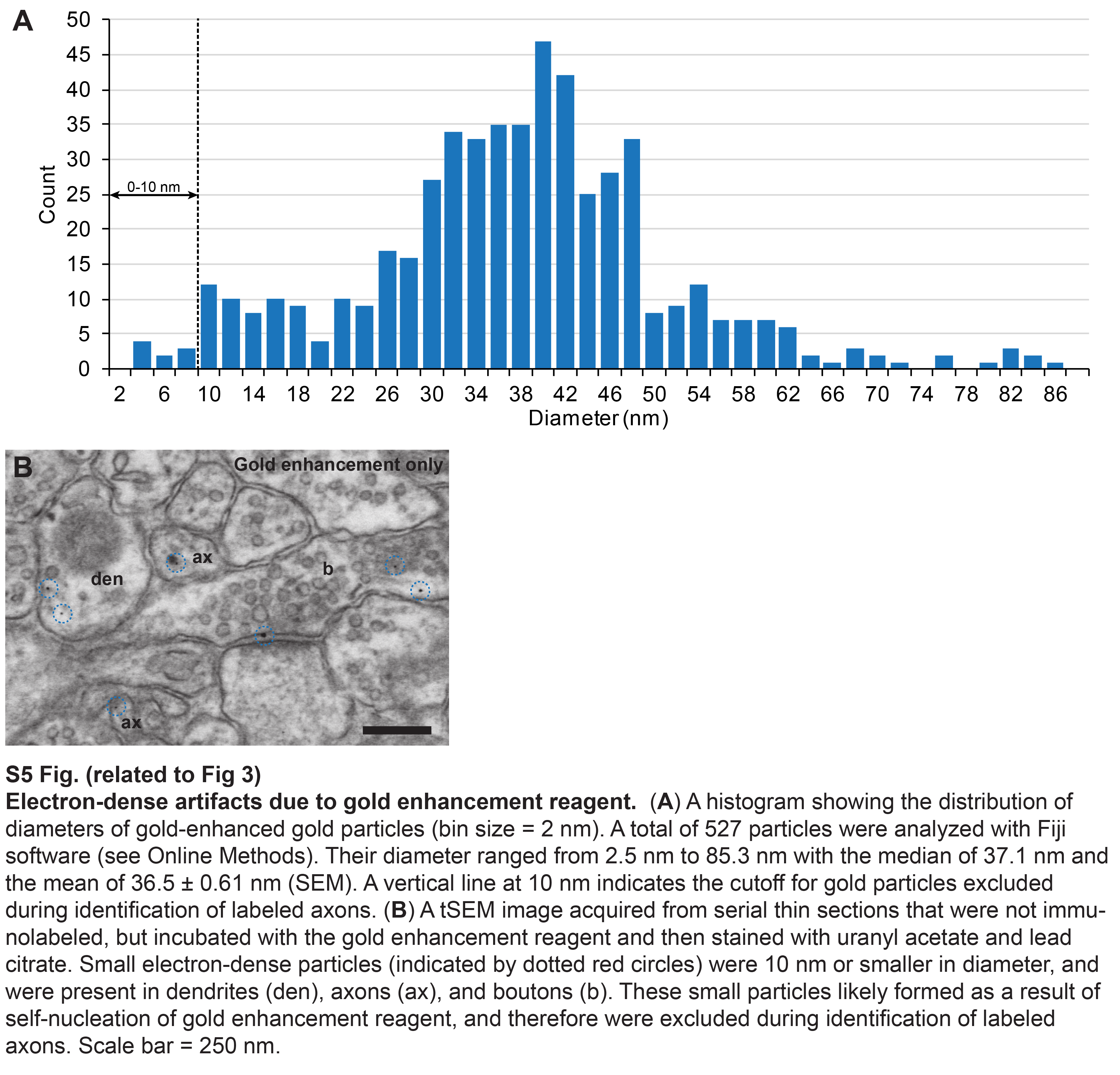
